# Probing Physical Intuition in Large Language Models through Partner-Dependent Mutation Prediction

**DOI:** 10.64898/2026.10.09.757856

**Authors:** Liu Cheng, Kaipeng Zheng, Weiran Huang, Hai-Feng Chen, Zhangsheng Yu

## Abstract

Structure-guided mutation design requires relating a proposed substitution to its molecular environment to anticipate its effects. Existing evaluations of large language models (LLMs) in protein engineering emphasize sequence-based prediction or tool-integrated workflows, leaving unclear whether LLMs themselves can infer mutation effects from structural evidence without dedicated predictors. We evaluate physical intuition—qualitative reasoning that connects structural observations with molecular interaction principles—using a benchmark that tests the same substitution in the same protein across different binding partners, with predictions based on local atomic coordinates and no dedicated mutation-effect predictors. In the tested comparisons, the best-performing LLM achieved group accuracy comparable to the evaluated energy predictors and Biomni, an LLM-driven tool workflow, while the methods differed in their prediction patterns. Most evaluated LLMs achieved higher group accuracy for larger affinity improvements. Structural interventions support selective dependence on mutation-site and partner information, while assessable structural claims largely agree with the supplied coordinates. These findings support direct LLM structural reasoning as a basis for developing rational mutation-design tools that connect molecular observations with partner-specific effects.

## 1 Introduction

Structure-guided protein design requires relating a proposed mutation to its molecular environment. Human experts routinely inspect local geometry and chemistry to identify interactions that a substitution may alter and to judge whether the surrounding structure can accommodate the change (Kortemme et al., 2004; Kortemme and Baker, 2002). These judgments are inherently context dependent. The same substitution can strengthen binding to one partner while weakening binding to another because packing, electrostatics, hydrogen bonding, and conformational accommodation differ across interfaces (Kortemme et al., 2004; Havranek and Harbury, 2003). This context dependence is central to specificity engineering: a useful judgment must connect structural evidence to partner-specific consequences rather than treat a substitution as intrinsically beneficial or detrimental.

Such reasoning is not limited to unaided human intuition. Classical structure-based modelling frameworks formalize many of the same physical and chemical considerations that guide expert judgment, encoding them in scoring functions, interaction terms, and conformational sampling procedures (Schymkowitz et al., 2005; Alford et al., 2017; Barlow et al., 2018). In this sense, rational protein design spans a continuum from qualitative structural inspection by experts to explicit computational models of molecular interactions. What these approaches share is that mutation assessment is grounded in structural evidence and in general principles governing molecular interactions (Selzer et al., 2000; Barderas et al., 2008). Deep-learning methods extend these capabilities to sequence design conditioned on backbone or atomic context (Dauparas et al., 2022; Hsu et al., 2022; Dauparas et al., 2025), as well as variant-effect prediction from sequence and evolutionary information (Meier et al., 2021; Frazer et al., 2021; Notin et al., 2022). Their utility is supported by experimental applications such as antibody affinity maturation (Hie et al., 2024), while benchmarks such as ProteinGym assess predictive performance across diverse experimental assays (Notin et al., 2023). Nevertheless, predictive advantages remain task dependent: FoldX and Rosetta Flex ddG remain competitive with specialized deep-learning predictors in challenging affinity-prediction settings (Cai et al., 2024). Predictive scores also do not necessarily provide a mechanistic account of how a substitution affects its molecular environment (Burgin, 2025).

General-purpose large language models (LLMs) are increasingly evaluated in tool-integrated protein-engineering and biomedical workflows (Ghafarollahi and Buehler, 2024; Swanson et al., 2025; Huang et al., 2026; Tan et al., 2026; Kim and Romero, 2026; Cai et al., 2026). In many such systems, the LLM plans analyses, invokes specialized modelling or prediction tools, reads their outputs, and uses those outputs to guide subsequent decisions (Xu et al., 2026; Teneggi et al., 2026). This mode of tool use can be highly effective, but end-to-end workflow performance does not isolate whether the LLM itself can make the kind of structure-based judgment that underlies rational design. A specialized component may already supply an estimate of the mutation effect (Cai et al., 2025), leaving the LLM primarily to interpret, communicate, or act on that prediction. PG-LLM evaluates tool-free variant ranking from protein sequences and assay descriptions, without providing protein structures (Arora et al., 2026). Although this setting tests whether LLMs can predict variant effects, it does not isolate whether they can reason about how the same substitution acts in different binding environments. Successful predictions may draw on prior knowledge of proteins and mutations or on sequence-based inference; they do not by themselves establish that a model can examine a particular protein–partner complex and connect its structural features to partner-specific consequences. This motivates a more direct question: can an LLM itself identify and integrate mutation-relevant structural evidence to make partner-dependent judgments without a dedicated predictor of the target effect?

To address these questions, we construct a benchmark that holds the focal protein and amino-acid substitution fixed across different binding partners, drawing on SKEMPI 2.0 and literature-supported systems (Jankauskaitė et al., 2019). This design tests whether predictions respond appropriately to the binding environment while the substitution itself remains unchanged. Each complex is presented independently, without revealing group membership, and a group is counted as correct only when every partner-specific outcome is predicted correctly. Models inspect structural coordinates with or without access to code execution for structural analysis, while dedicated energetic and mutation-effect predictors are excluded. Structural interventions, audits of model-generated claims, and mechanistic case studies then examine the evidence underlying these judgments and the limits of its interpretation.

Our results show that direct structural reasoning by general-purpose LLMs can achieve group-level prediction accuracy comparable to established physical predictors across both datasets, without relying on dedicated predictors to supply mutation-effect judgments. The best-performing LLM also matches or exceeds the tool-integrated workflow Biomni (Huang et al., 2026) in the evaluated comparisons. Similar accuracies nevertheless conceal different prediction patterns: LLMs exhibit distinct precision–recall profiles relative to the physical predictors, and the matched LLM–Biomni comparison reveals partially different sets of correctly resolved groups. Success also depends on effect magnitude. Most evaluated LLMs and both physical predictors recover more complete partner contrasts in the larger-improvement subset, consistent with the expectation that more pronounced affinity improvements are easier to predict than subtle changes. These judgments also show selective dependence on the supplied structural evidence: mutation-site masking changes predictions more often than distant masking, removing partner information or atomic coordinates reduces accuracy, and auditable structural claims largely agree with the input structures. Together, these findings provide systematic evidence that LLMs can use three-dimensional structural information to make partner-dependent mutation judgments, with performance comparable to established methods in the tested settings, distinct prediction patterns, and greater success for larger affinity improvements.

## 2 Benchmark and evaluation protocol

We organize the benchmark into mutation groups that hold the focal protein and amino-acid substitution fixed across different binding partners (Fig. 1). The benchmark comprises 60 distinct groups and 126 mutation–partner observations: 56 groups include both affinity-enhancing and affinity-weakening observations, while four contain only affinity-enhancing observations. Dataset 1 is curated from SKEMPI 2.0, whereas Dataset 2 is assembled through a PDB-wide search followed by manual verification of structures and experimental mutation effects reported in the literature. Appendix A provides detailed curation and input-processing procedures.

**Figure 1:**
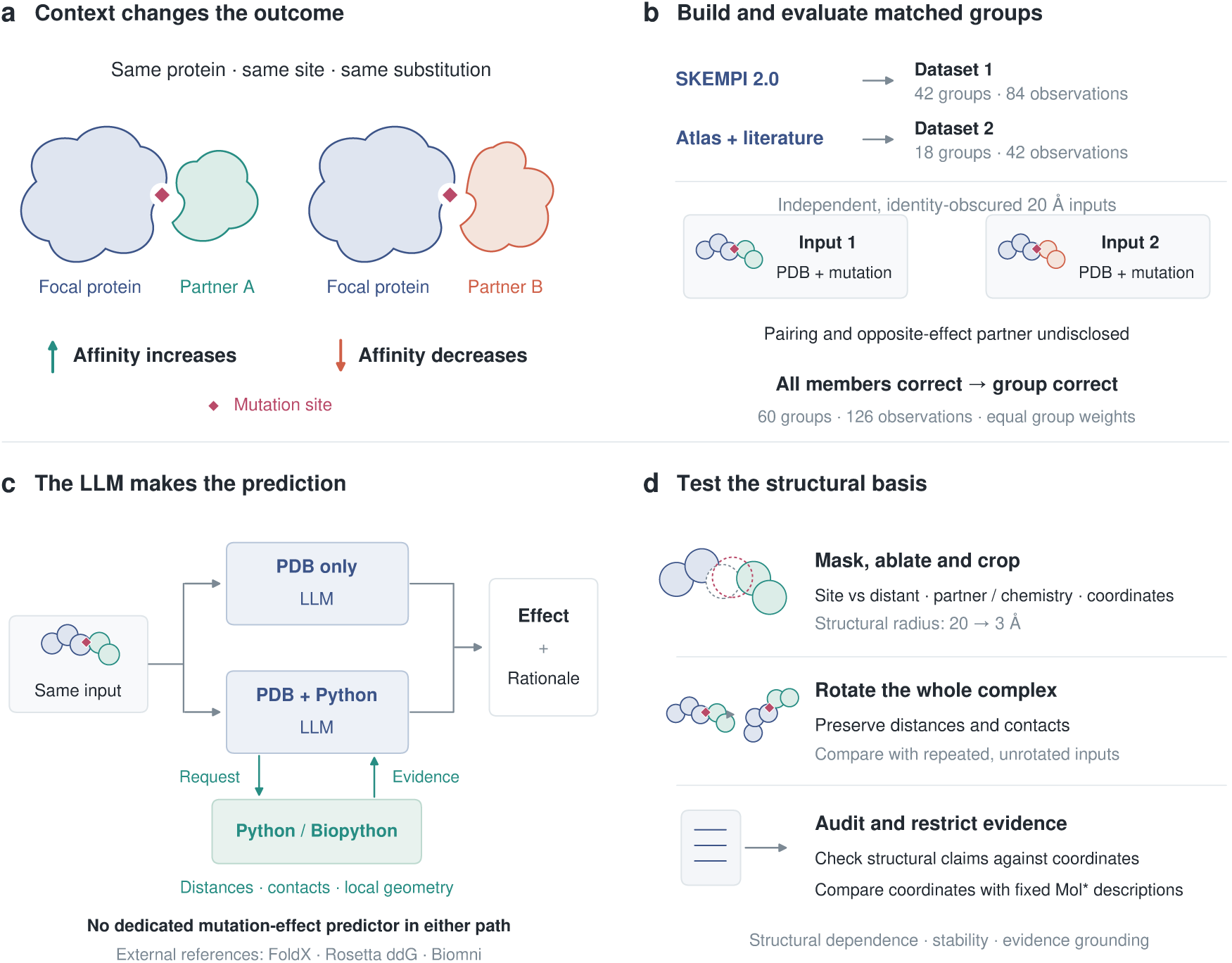
Benchmark design and evaluation. a, The same substitution in a fixed protein can have opposite effects across binding partners. b, Two curated datasets comprise 60 groups and 126 mutation–partner observations. Complexes are assessed independently; a group is correct only when every partner-specific outcome is predicted correctly. c, Models predict mutation effects from structural coordinates with or without code execution, without dedicated mutation-effect predictors. d, Structural interventions, coordinate rotations, claim audits and comparisons with fixed structural descriptions probe the evidence underlying predictions.

### 2.1 Dataset curation

Dataset 1 retains single substitutions at binding interfaces with measured wild-type (WT) and mutant dissociation constants from SKEMPI 2.0 (Jankauskaitė et al., 2019). Effects are defined as 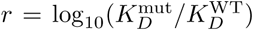, with negative values indicating enhancement. Repeated measurements are represented by their median log ratio. We retained only groups with exactly two eligible partners and opposite nonzero mutation effects, without imposing an additional effect-magnitude threshold. This yielded 42 groups and 84 observations across 17 WT sequences. All mutation sites lie within 5 Å of partner-protein heavy atoms.

To complement SKEMPI 2.0 with additional experimental evidence, particularly from studies published after its release, we assembled Dataset 2 through a PDB-wide search (Berman et al., 2000) and manual literature curation. Candidate complexes were checked against the original publications to verify structural correspondence and experimentally reported mutation effects. After excluding Dataset 1 overlap, it contains 18 groups and 42 observations: 14 opposite-effect groups and four all-enhancing groups. The latter assess recovery of beneficial mutations across partners, rather than effect reversal. Their inclusion was motivated by observed negative-prediction tendencies and the affinity-improvement objective. Dataset 2 includes groups with more than two observations and one protein–small-molecule group; it does not estimate the natural prevalence of beneficial mutations.

### 2.2 Structural inputs and inference conditions

For each complex, the model independently received a mutation specification and structural coordinates from a 20 Å neighbourhood around the mutation site. Molecular identifiers and group membership were withheld; renumbering and record permutation reduced direct identity cues without guaranteeing that identity was unrecoverable (Appendix A.5). An identity-recovery check with DeepSeek V4.1 Flash (hereafter, DeepSeek Flash) reported 16/84 successes for processed PDBs versus 76/84 for reference inputs (Appendix A.6, including reporting qualifications).

Models received identical coordinate records in two conditions: PDB only, without external tools or code execution, and PDB + Python, with access to a Python environment and basic libraries such as Biopython (Cock et al., 2009) for structural inspection. Dedicated energetic and mutation-effect predictors were excluded; the LLM selected and interpreted evidence to predict the effect direction.

### 2.3 Evaluation metrics and comparisons

The primary endpoint is group accuracy, with equal weight per group and every member required to be correct. LLM method comparisons use majority-vote consensus from three runs. Observation-level precision and recall describe enhancement predictions; task accuracy provides an additional descriptive measure.

For each Dataset 1 pair, its enhancing member defines 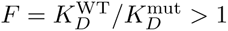. We compare 21 larger-improvement groups (*F ≥* 2) with 21 smaller-improvement groups (1 *< F <* 2). Both strata contain one enhancing and one weakening observation per group.

We compare direct LLM judgments with structure-based energy predictors, FoldX and Rosetta ddG (Schymkowitz et al., 2005; Barlow et al., 2018), and an LLM-driven tool-integrated workflow, Biomni (Huang et al., 2026). Physical predictors use full structures and exclude the Dataset 2 protein–small-molecule group, leaving 17 groups and 40 observations.

## 3 LLM performance on partner-dependent mutation prediction

### 3.1 Larger affinity improvements are associated with higher group accuracy

Overall group accuracy pools contrasts with different effect magnitudes. We compare prediction performance across two disjoint Dataset 1 strata defined by larger and smaller affinity improvements, each containing one affinity-enhancing and one affinity-weakening observation per group. Three of the four LLMs evaluated with access to code execution and both physical predictors achieved higher group accuracy in the larger-improvement stratum (Fig. 2; Table 1). GLM-5.3 Flash resolved 7 of 21 groups (33.33%), compared with 1 of 21 (4.76%) in the smaller-improvement stratum. GLM-5.3 and DeepSeek Flash likewise achieved higher group accuracy in the larger-improvement stratum. FoldX and Rosetta ddG resolved seven versus zero and eight versus two groups, respectively. Qwen3.8 Flash was the exception, resolving two groups in each stratum.

**Figure 2:**
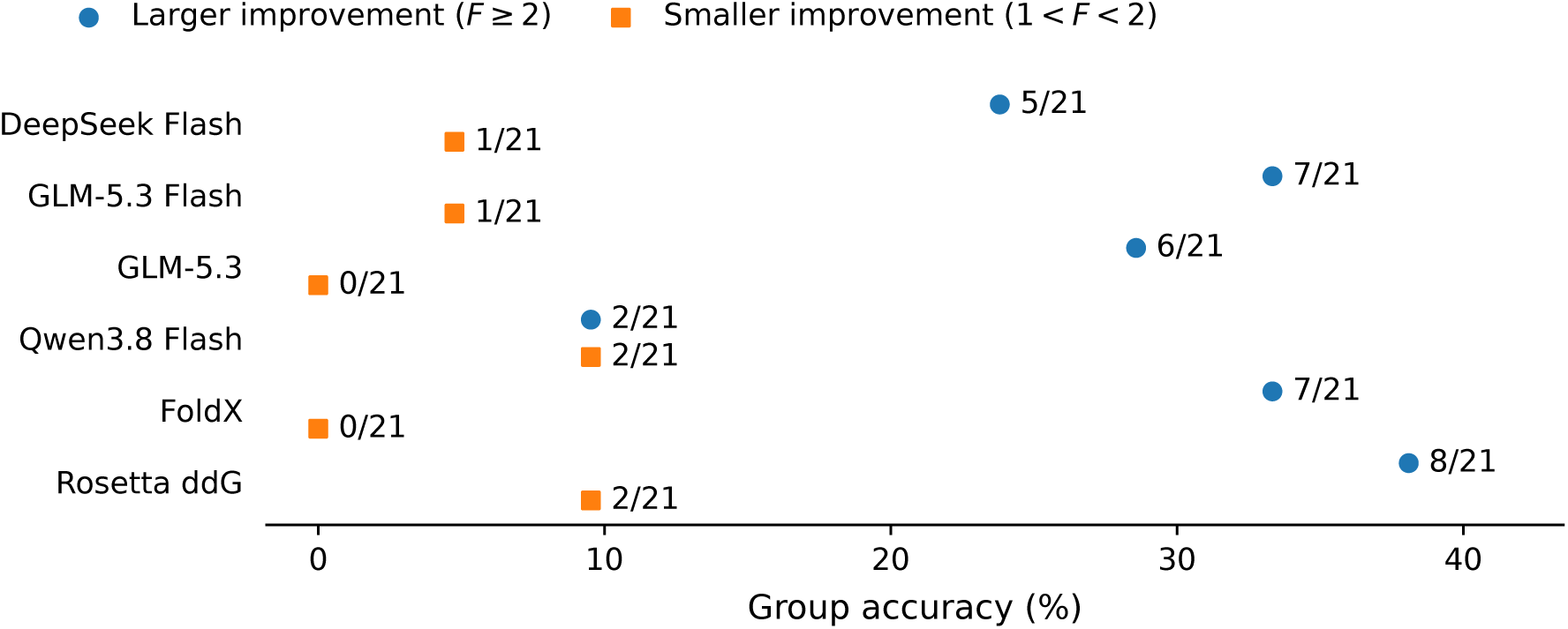
Most methods achieve higher group accuracy for larger affinity improvements. The two Dataset 1 strata each contain 21 enhancing–weakening pairs, classified by 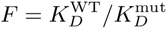 for the enhancing member (*F ≥* 2 versus 1 *< F <* 2). Both partner-specific predictions must be correct; labels show correct/total groups. LLM results use code execution and majority-vote consensus from three runs.

**Table 1:** Group accuracy across datasets and affinity-improvement strata.

| Method | Dataset 1 |  |  | Dataset 2 |
| --- | --- | --- | --- | --- |
| | All<br>42 groups | Larger improvement<br>$F \geq 2$: 21 groups | Smaller improvement<br>$1 < F < 2$: 21 groups | All<br>18 groups |
| *LLMs with access to code execution* |  |  |  |  |
| DeepSeek Flash | 14.29 (6/42) | 23.81 (5/21) | 4.76 (1/21) | 11.11 (2/18) |
| GLM-5.3 Flash | 19.05 (8/42) | 33.33 (7/21) | 4.76 (1/21) | 16.67 (3/18) |
| GLM-5.3 | 14.29 (6/42) | 28.57 (6/21) | 0.00 (0/21) | 11.11 (2/18) |
| Qwen3.8 Flash | 9.52 (4/42) | 9.52 (2/21) | 9.52 (2/21) | 0.00 (0/18) |
| *Physical predictors* |  |  |  |  |
| FoldX | 16.67 (7/42) | 33.33 (7/21) | 0.00 (0/21) | 11.76 (2/17)<sup>†</sup> |
| Rosetta ddG | 23.81 (10/42) | 38.10 (8/21) | 9.52 (2/21) | 5.88 (1/17)<sup>†</sup> |
| *Agentic workflow* |  |  |  |  |
| Biomni | — | 33.33 (7/21) | — | 5.56 (1/18) |

Values are percentages (correct/total groups); every partner-specific prediction must be correct. LLM results use code execution and majority-vote consensus from three runs. $F = K_D^{\text{WT}}/K_D^{\text{mut}}$ refers to the enhancing member of each Dataset 1 group.
<sup>†</sup> Physical predictors exclude the Dataset 2 protein–small-molecule group, leaving 17 groups.

These differences concern complete opposite-effect pairs, not just identification of the enhancing member. The higher accuracy for larger improvements across most evaluated LLMs and both physical predictors is consistent with the expectation that more pronounced mutation effects are easier to predict than subtler changes.

### 3.2 Comparable group accuracy with distinct prediction patterns

With access to code execution, the best-performing LLM, GLM-5.3 Flash, achieved group accuracy comparable to that of the evaluated physical predictors (Table 1). On Dataset 1, GLM-5.3 Flash resolved eight groups, compared with seven for FoldX and ten for Rosetta ddG. LLMs and physical predictors nevertheless show distinct precision–recall profiles (Fig. 3a): several LLMs make fewer but more precise enhancement predictions. DeepSeek Flash achieved 91.67% precision at 16.18% recall across both datasets, whereas the physical predictors recovered more enhancing observations.

**Figure 3:**
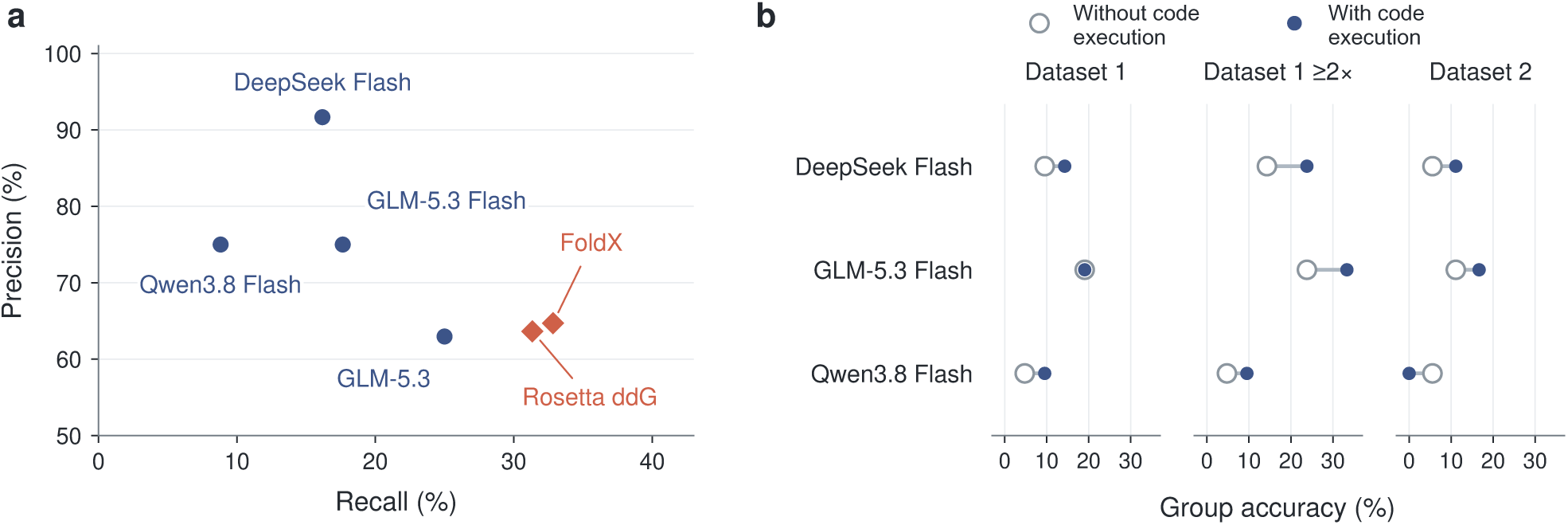
Prediction patterns and the contribution of code execution. a, Precision and recall for affinity enhancement pooled across both datasets. LLMs with code execution (circles) use 126 observations; energy predictors (diamonds) use 124, excluding the protein–small-molecule group. b, Group accuracy with (filled circles) and without (open circles) code execution on Dataset 1, its larger-improvement subset, and Dataset 2 (42, 21 and 18 groups, respectively).

On the matched 21-group larger-improvement subset, GLM-5.3 Flash and Biomni each resolved seven groups (33.33%). Five correct groups were shared, while each method resolved two missed by the other. Although absolute group accuracy remains modest, the best-performing LLM achieves results comparable to the evaluated energy predictors and Biomni. Despite similar overall accuracy, the methods differ in their precision–recall profiles and in which mutation groups they predict correctly.

### 3.3 LLMs can correctly predict partner-dependent effects without code execution

On Dataset 1, GLM-5.3 Flash resolved the same number of groups with and without access to code execution; DeepSeek Flash and Qwen3.8 Flash each resolved two fewer without it (Fig. 3b). Access to code execution provided additional gains on the larger-improvement subset, but effects on Dataset 2 varied. Without code execution, DeepSeek Flash and Qwen3.8 Flash succeeded only on the same all-enhancing group, whereas GLM-5.3 Flash resolved two opposite-effect groups. Thus, code execution is not necessary for every successful partner-dependent judgment, but the results do not establish equivalence between tool conditions. Appendix A.4 separates Dataset 2 successes by group composition.

### 3.4 Prediction success spans multiple protein systems

With access to code execution, DeepSeek Flash, GLM-5.3 Flash and Qwen3.8 Flash resolved Dataset 1 groups across four, five and four WT protein systems, respectively, collectively covering seven of the 17 systems (Appendix C, Fig. S7). Successes also span multiple systems without code execution. Coverage is uneven: BPTI accounts for half of the successes of GLM-5.3 Flash with access to code execution and of Rosetta ddG.

## 4 LLM judgments depend on structural evidence

We next examine whether LLM predictions depend on the structural evidence supplied for each complex. We selectively mask or remove structural information and measure how the predictions change.

### 4.1 Predictions depend on mutation-site and partner information

Mutation-site masking changed predictions more often than distant side-chain masking. With access to code execution, mutation-site masking changed at least one consensus prediction in 42.06% of Dataset 1 groups, averaged across DeepSeek Flash, GLM-5.3 Flash and Qwen3.8 Flash, compared with 10.32% for distant side-chain masking (Fig. 4a). Changes were determined by comparing the consensus predictions for masked and intact inputs. Mutation-site masking changed predictions more often than distant masking for all three models. Without code execution, mutation-site masking changed at least one consensus prediction in a mean 19.05% of groups across the same models. The stronger response to mutation-site masking indicates selective dependence on local structural information.

**Figure 4:**
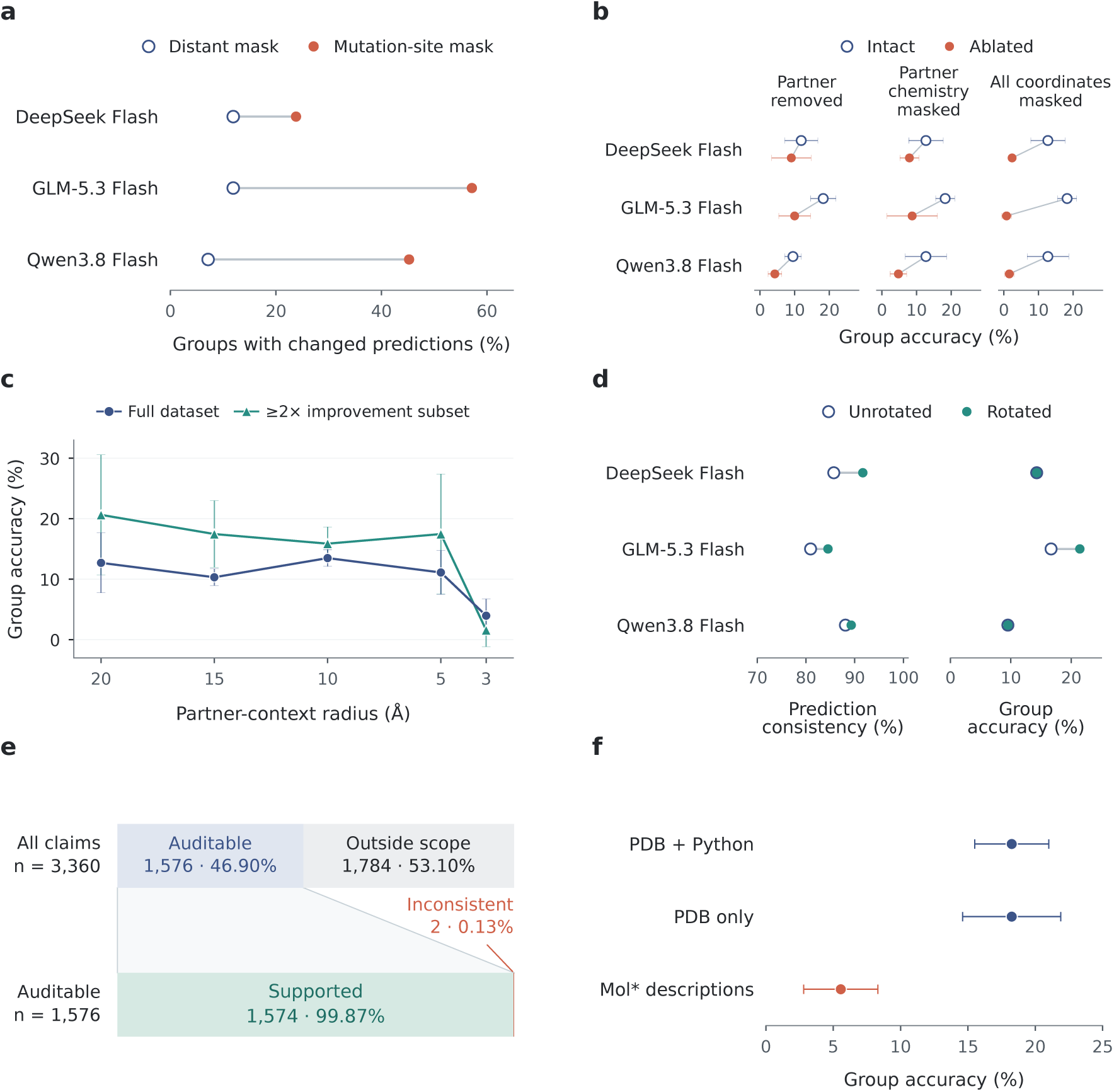
Dependence of mutation judgments on structural evidence. Results use Dataset 1 (42 groups, 84 observations). a, Groups with at least one changed consensus prediction after masking, with code execution. b, Group accuracy after partner removal, partner-chemistry masking or coordinate removal, each paired with its intact control. c, DeepSeek Flash accuracy across partner-context radii, with the focal-protein input fixed at 20 Å. d, Consistency and accuracy under rotated or unchanged coordinates; consistency requires identical predictions across three runs within each condition. e, Scope and coordinate agreement of audited structural claims. f, GLM-5.3 Flash accuracy with coordinate inputs or fixed Mol*-derived descriptions.

Each ablation was compared with its corresponding intact-input control. Removing the partner chain lowered mean group accuracy across the three models from 13.22% to 7.78% (Fig. 4b). In a separate experiment, retaining the partner backbone while masking its side chains and residue identities reduced mean group accuracy from 14.55% to 7.14%. Both interventions reduced prediction accuracy, indicating that the models used information about the binding partner.

Removing all atomic coordinates while retaining residue identities and other non-geometric information reduced mean group accuracy from 14.55% to 1.59%. Mean enhancement recall fell from 17.20% to 2.38%, while specificity remained high. Without coordinates, the models predominantly predicted affinity weakening and rarely recovered enhancing mutations.

For DeepSeek Flash, we varied the partner-context radius from 20 Å to 15, 10, 5 and 3 Å, while keeping the focal-protein portion of the original 20 Å input unchanged. We used three runs per condition (Fig. 4c). Full-dataset mean group accuracy ranged from 10.32% to 13.49% across the tested 5–20 Å radii. At 3 Å, it fell to 3.97%, compared with 12.70% at 20 Å. In the larger-improvement subset, mean group accuracy fell from 20.63% at 20 Å to 1.59% at 3 Å. The largest observed drop occurred when the partner-context radius was reduced to 3 Å.

### 4.2 Tested coordinate rotations do not reduce prediction consistency or accuracy

We applied three independent rotations to each complex as a rigid body, preserving relative atomic geometry, and compared predictions with three repeated runs using unchanged coordinates. Within each condition, consistency was defined as the proportion of mutation–partner observations receiving the same prediction in all three runs. Averaged across the three models, consistency was 88.49% with rotated coordinates and 84.92% with unchanged coordinates. Consistency was higher for all three models (Fig. 4d). Group accuracy, averaged across the three models, was 15.08% with rotated coordinates and 13.49% with unchanged coordinates; the corresponding larger-improvement-subset values were 23.81% and 17.46%. These comparisons assess consistency within each condition and aggregate accuracy; they do not establish that each observation receives the same prediction before and after rotation.

### 4.3 Assessable structural claims largely agree with the supplied coordinates

We audited 3,360 structural claims from 1,344 Dataset 1 predictions, including contacts, distances and residue-level spatial relationships (Fig. 4e). Of these, 1,576 claims (46.90%) were within the predefined audit scope. Coordinate-based checking with Mol*, followed by manual review of flagged conflicts, supported 1,574 of these claims (99.87%) and identified two as inconsistent with the supplied coordinates. The remaining 1,784 claims concerned properties outside the audit scope, including solvent accessibility and qualitative pocket geometry. The assessable structural observations therefore largely agreed with the supplied coordinates. This agreement does not validate unassessed claims or establish that the cited interactions explain the experimentally observed mutation effects. The audit procedure is detailed in Appendix D.

### 4.4 Fixed structural descriptions yield lower accuracy than coordinate inputs

We replaced coordinates with fixed structural descriptions generated using Mol*, covering predefined contacts, distances and local-environment features for the same neighbourhoods (Sehnal et al., 2021). GLM-5.3 Flash achieved 5.56% *±* 2.75% group accuracy with descriptions, versus 18.25% *±* 3.64% with PDB alone and 18.25% *±* 2.75% with PDB + Python (three-run means *±* sample standard deviations; Fig. 4f). Group accuracy was lower with the fixed descriptions than with either coordinate-input condition. This comparison does not determine whether the gap reflects differences in evidence coverage, presentation format, or opportunities for the model to select mutation-relevant structural features.

These results show that LLM predictions depend selectively on structural information and that assessable structural observations largely agree with the supplied coordinates. We next compare model explanations with experimentally supported mutation mechanisms in selected case studies.

## 5 Case studies of LLM structural reasoning

These cases reveal a progressive picture of LLM-based mutation-effect reasoning, from recognizing partner-dependent determinants to evaluating mutant-state mechanisms and mechanistic validity.

### Partner-dependent interpretation

We first examined whether LLMs can recognize that mutation effects are determined by the local binding environment rather than by the mutation itself. The ShPI-1 K13L case provides a direct test because the same substitution produces opposite effects against different protease partners (García-Fernández et al., 2015). GLM-5.3 Flash correctly predicted that K13L improves *α*-chymotrypsin inhibition but weakens trypsin inhibition. For *α*-chymotrypsin, the model identified the unfavorable energetic consequence of a buried lysine lacking sufficient electrostatic compensation and proposed that leucine could provide more favorable hydrophobic packing (Fig. 5a). For trypsin, GLM-5.3 Flash similarly identified K13 as a deeply buried interface anchor stabilized by electrostatic interactions and hydrogen-bonding networks; therefore, K13L would disrupt this polar stabilization network (Fig. 5b). Thus, LLMs can capture partner-dependent structural determinants when the energetic consequences are strongly encoded by the local interaction environment.

**Figure 5:**
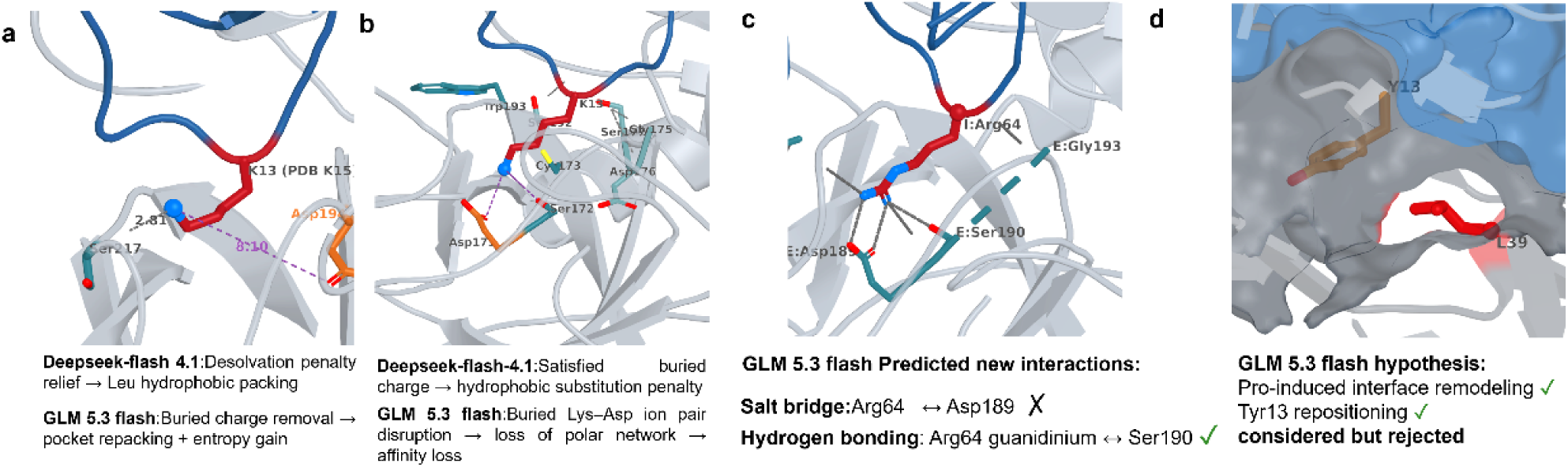
Structural interpretation and mutant-state hypotheses in selected mutation cases. a,b, ShPI-1 K13L in complexes with *α*-chymotrypsin (3T62) and trypsin (3M7Q), respectively. DeepSeek Flash and GLM-5.3 Flash interpret the distinct pocket environments as favouring hydrophobic substitution in the former and loss of polar stabilization in the latter. The structures show the starting Lys residue; literature Lys13 corresponds to PDB Lys15 in a. c, BbKI F64R in complex with KLK4. GLM-5.3 Flash proposed Arg64-mediated contacts from the Phe64 input structure 7JQV; the displayed Arg64 structure 7JOD is an experimental comparison, not the model input or an LLM-generated structure. The Ser190 contact is supported, whereas the proposed direct bidentate Arg64–Asp189 salt bridge requires assessment of atom geometry and alternate conformations. The cross marking this interaction should not be taken as ruling out all Arg–Asp contacts. d, EVA-A L39P in complex with CCL7, shown in the WT input structure 7SCU. GLM-5.3 Flash considered favourable interface remodelling and Tyr13 repositioning but ultimately predicted no improvement, emphasizing losses from the starting-state interaction network. The proposed repositioning is directionally consistent with the experimental L39P comparison 8FK8. Structural views show experimental coordinates; highlighted residues and contact lines locate the interactions discussed, without assigning their energetic contributions. Detailed comparisons are provided in Appendix B.

### Mutant-state interaction prediction

We next examined whether LLMs can infer mutation-induced interaction changes from WT structures. The BbKI F64R cases were analyzed in two distinct complexes involving bovine trypsin and human kallikrein 4 (KLK4) (Li et al., 2021). The results show that LLMs can propose relevant structural changes after mutation, including the formation of Arg64-mediated interactions, water displacement, and local interaction-network remodeling. Although several predicted Arg64-mediated contacts were partially supported by experimentally determined mutant structures, some specific interactions were overestimated (Fig. 5c). Notably, DeepSeek Flash and GLM-5.3 Flash proposed similar mutant-state hypotheses, but reached different final predictions because they assigned different weights to the energetic contributions inferred from structural geometry. These results indicate that LLMs can generate plausible mutant-state hypotheses, but remain limited in translating local interaction changes into overall binding free-energy effects. More broadly, uncertainty arises from two major challenges: extrapolating unobserved mutant conformations from known WT structures and converting structural changes into energetic consequences for decision-making.

### Difficulty in evaluating indirect structural effects

The EVA-A Y44A and L39P cases (Devkota et al., 2023) revealed another limitation: in these cases, affinity changes were mediated by indirect structural effects rather than by direct interactions formed by the mutated residues, and therefore the evaluated LLMs failed to correctly predict these cases (Fig. 5d). Detailed analysis of the reasoning trajectories showed that the major difficulty was not the inability to propose plausible mechanisms or mechanisms consistent with the literature, but rather an asymmetric weighting between WT structural evidence and mutant-state hypotheses. Notably, these models did generate hypotheses consistent with the experimentally observed mechanisms, including ligand relaxation and interface repositioning. However, because the models could only observe static experimentally determined structures, these hypotheses were rapidly self-rejected during the reasoning process due to the dominance of static structural evidence and WT-based interactions.

### Correct prediction with incorrect reasoning

Finally, the BLIP E73W case demonstrates that correct prediction of the affinity-effect direction does not necessarily indicate mechanistic understanding (Rivera et al., 2025). Although GLM-5.3 Flash correctly predicted the affinity enhancement and identified several valid wild-type (WT) structural features, the proposed mechanistic explanation was not consistent with the mechanism described in the literature. The model attributed the affinity improvement mainly to the removal of the unfavorable energetic penalty associated with the buried Glu residue, water displacement, and aromatic repacking. However, the mutant structure revealed a distinct interaction mode involving packing between Trp73 and Tyr105 and stabilization of an alternative partner conformation. This case shares a similar failure mode with the EVA-A cases (Y44A and L39P). When LLMs lack relevant prior knowledge or the ability to reason beyond static geometric information, they face challenges in more complex mutation-effect interpretation problems.

Together, these cases suggest that LLMs possess the ability to extract chemically meaningful structural determinants from observed protein geometries and generate experimentally testable mutation hypotheses. However, when performing mutation-effect reasoning from WT structures, their major limitation lies in extrapolating beyond directly observed structural information, including mutant-state adaptation, conformational changes, and the energetic consequences of local interaction remodeling.

## 6 Discussion

This study provides evidence that general-purpose LLMs can make partner-dependent mutation judgments from structural information without a dedicated predictor supplying the target effect. The strongest evaluated model achieved group accuracy comparable to the tested energy predictors and the tool-integrated workflow Biomni, while showing different prediction patterns. Prediction success was also associated with effect magnitude: most evaluated LLMs achieved higher group accuracy for larger affinity improvements. The relevant capability is to connect a substitution with the molecular environment of a particular complex and reach a partner-specific conclusion. Absolute accuracies nevertheless remain modest, with substantial room for improvement in reliability and coverage.

The structural interventions help establish that this capacity involves information supplied in the coordinates. Predictions responded more strongly to mutation-site masking than to distant masking, and accuracy decreased when partner information or atomic coordinates were removed. The audit independently showed that assessable structural claims largely agreed with the supplied coordinates. These findings jointly support selective use of structural evidence, although such use may coexist with learned knowledge about proteins, mutations and molecular interactions. Here, physical intuition denotes qualitative inference connecting structural observations with principles of molecular interaction; it makes no assumption about human-like internal reasoning. Successful predictions without code execution further show that external computation is not necessary for every correct judgment. However, the results do not identify a unique internal representation or reasoning process through which the models use structural information.

The case studies suggest that a major challenge lies in moving from the observed structure to plausible mutant states and their energetic consequences. A starting structure provides evidence about existing contacts and local geometry, but mutation assessment requires anticipating which interactions will persist, disappear or be replaced after substitution. In selected cases, model explanations proposed rearrangements consistent with experimental observations but ultimately favoured conclusions based on the starting interaction network. Generating a plausible mutant-state hypothesis was therefore insufficient to secure a correct prediction. These patterns suggest difficulties in comparing alternative structural hypotheses and assigning appropriate weight to their consequences. Because the analysis concerns expressed explanations, it cannot establish how those explanations relate to internal decision processes. It does, however, identify concrete targets for evaluation: predicting structural adaptation, comparing competing hypotheses and integrating their consequences into an overall affinity judgment.

More broadly, these findings suggest a route toward a new generation of rational protein-design tools. Frameworks such as FoldX and Rosetta translate accumulated biophysical knowledge into explicit energy terms, calibrated parameters and structural modelling procedures (Schymkowitz et al., 2005; Alford et al., 2017). The capacity of LLMs to connect structural observations with partner-dependent mutation effects raises the possibility of extending this approach through models that can formulate, combine and revise structural reasoning strategies. Coupled with systematic computational and experimental feedback, such models could help identify limitations in existing approximations, propose new scoring heuristics and develop improved methods for mutation assessment. The longer-term opportunity is to translate this reasoning capacity into computational procedures that can be inspected, tested and refined, potentially yielding more effective tools for rational mutation design. Realizing this potential will require showing that the resulting methods improve prediction and design outcomes on independent systems and in prospective experiments.

## 7 Conclusion

We show that general-purpose LLMs can use supplied structural evidence to make partner-dependent mutation judgments without dedicated mutation-effect predictors. The best-performing model achieved group accuracy comparable to the evaluated energy predictors and Biomni in the tested comparisons, while the methods differed in their prediction patterns. Most evaluated LLMs achieved higher group accuracy for larger affinity improvements, and successful judgments also occurred without code execution. Structural interventions and claim audits support selective dependence on mutation-relevant information and the accuracy of assessable structural observations. Overall predictive accuracy nevertheless remains modest, and valid structural observations do not ensure correct mutation-effect predictions or mechanistic explanations. These findings provide a basis for developing a new generation of rational mutation-design tools that incorporate LLM structural reasoning, with improved assessment of mutant states and their energetic consequences as important directions for further development.

## Acknowledgements

This work was supported by Shanghai Municipal Science and Technology Major Project, partially by SJTU Kunpeng & Ascend Center of Excellence, the Center for HPC at Shanghai Jiao Tong University, and the National Key Research and Development Program of China (2025YFA0921000 and 2023YFF1205102), National Natural Science Foundation of China (ID: 12171318 and 32571435), Shanghai Science and Technology Commission (ID: 24JS2810200, 21ZR1436300, 23XD1401900, 23DZ2290600), the Fundamental Research Funds for the Central Universities (YG2023LC03) and Medical Engineering Cross Fund of Shanghai Jiao Tong University (ID: YG2023ZD21).

## A Additional benchmark curation and input-processing details

### A.1 Dataset 1: filtering, identity mapping and aggregation

Initial deduplication reduced 7,085 SKEMPI records to 6,794 measurement records before quality filtering and aggregation of repeated experimental measurements. We required finite, positive WT and mutant dissociation constants and excluded censored affinities. Original and cleaned single-substitution annotations had to agree, and the mutated chain had to resolve unambiguously to one side of the complex and a unique full-length WT sequence. WT residue identity was checked before and after sequence mapping. Mutation groups were defined by the exact WT sequence and mapped substitution.

Partner identity was the sorted, deduplicated set of complete partner-chain sequences, independent of protein names or PDB numbering. Full-length UniProt WT sequences were used when all partner chains mapped unambiguously; otherwise, the entire partner side was represented by complete PDB polymer sequences. This definition does not distinguish repeated copies of an identical chain. Repeated measurements for the same complex, WT sequence, substitution and partner were consolidated using the median of individual log ratios, rather than the log ratio of separately aggregated WT and mutant affinities. Experimental conditions and publications were retained for auditing but did not partition this aggregation.

Interface filtering used the first PDB model and the highest-occupancy alternate location for each atom name. The mutated WT residue had to lie within 5 Å of a non-water heavy atom on a partner chain. Groups with more than two eligible partner identities were excluded from Dataset 1 rather than expanded into pairwise comparisons. If multiple eligible complex records represented a partner, the record supported by the most original measurements was selected; ties were broken lexicographically by complex identifier. Selected partner effects had to have strictly opposite signs; zero effects were excluded without imposing an additional effect-magnitude threshold. The resulting 42 groups spanned 17 WT sequences and 32 PDB entries. All 84 final observations passed a second 5 Å interface check using only partner-protein heavy atoms and were supported by 122 unique source measurements.

**Table S1:** SKEMPI curation stages.

| Stage | Retained records or groups |
| --- | --- |
| Raw SKEMPI records | 7,085 |
| After initial deduplication | 6,794 |
| After record-level quality filtering | 4,025 |
| After repeated-measurement aggregation | 3,396 evidence records |
| After interface filtering | 2,445 evidence records |
| Groups with exactly two partner identities | 259 groups |
| Selected groups with opposite effects | 42 groups / 84 observations |

### A.2 Effect-magnitude stratification

The enhancing member of each Dataset 1 pair defines 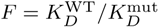. The larger-improvement stratum contains 21 groups with *F ≥* 2; the smaller-improvement stratum contains 21 with 1 *< F <* 2. Both retain one enhancing and one weakening observation per group. The smaller-improvement stratum is an internal comparison for subtler experimental effects, not a discarded set. No additional magnitude threshold applies to the weakening member, so neither stratum requires both members to have effects of a specified magnitude.

Stratum results reuse the full-dataset predictions. The source summary labels the FoldX subset results as FoldX5.

The comparison is descriptive and uses two bins within the same benchmark. It does not establish a continuous effect–accuracy relationship, matching of protein systems or mutation types between strata, or independently measured differences in structural salience. No statistical-significance or prediction-bias-adjustment claim is inferred from the stratum counts alone.

### A.3 Dataset 2: candidate generation and literature curation

The PDB-wide atlas began with 259,129 identifiers and linked focal proteins to distinct partners using biological assemblies, sequence mappings and direct-contact evidence. PDBe biological-assembly and molecular-entity annotations were combined with UniProt chain mappings. Co-assembly associations required direct-contact support from PDBe PISA or ligand binding-site annotations. Focal proteins required confirmed non-self contacts in at least two distinct PDB entries with at least two distinct raw partner identities. This generated 33,321 structural candidates without imposing partner-class, resolution or minimum interface-size thresholds at this stage.

Candidates were manually assessed for a fixed protein and substitution, identifiable partner complexes and literature-supported affinity effects. Structural contact alone was insufficient for inclusion. Despite extensive structural and literature searches, experimentally characterized multi-partner mutation groups were scarce. Fourteen groups contained opposite-sign effects. We additionally included four groups in which all measured effects were affinity-enhancing, motivated by the models’ observed tendency towards negative predictions and the practical goal of identifying beneficial mutations. These groups assess recovery of improvements across partners rather than opposite-effect discrimination. Groups containing only affinity-weakening observations were not included because they did not address this affinity-improvement objective. This selection reflects the evaluation objective, not the natural prevalence of beneficial mutations.

The 14 opposite-effect groups comprised ten groups with one enhancing and one weakening observation, two with two enhancing and one weakening observation, and two with two of each. Each of the four all-enhancing groups contained two observations. The final 18 groups comprised 42 observations (26 enhancing and 16 weakening), did not overlap Dataset 1, and included groups with more than two observations and one protein–small-molecule group. Excluding the latter left 17 groups and 40 observations (25 enhancing and 15 weakening) for physical-predictor comparisons. All groups were scored using the same requirement that every member be predicted correctly.

### A.4 Dataset 2 successes by group composition

The four all-enhancing groups each contain two positive observations. Only group 00401fc010988ad75ce4 was correctly resolved, by DeepSeek Flash and Qwen3.8 Flash with PDB-only input. No other evaluated method correctly resolved this group, and no method correctly resolved any of the other three all-enhancing groups. Correctly predicting only one member does not count as a correct group.

Accordingly, all correct Dataset 2 groups for the Python-assisted LLMs came from the 14 opposite-effect groups: DeepSeek Flash resolved two, GLM-5.3 Flash three, GLM-5.3 two, and Qwen3.8 Flash none. Biomni resolved one opposite-effect group. FoldX and Rosetta ddG resolved two and one, respectively, among the 13 opposite-effect groups available to physical predictors after excluding the protein–small-molecule group. Without Python, DeepSeek Flash and Qwen3.8 Flash each resolved one all-enhancing group and no opposite-effect groups, whereas GLM-5.3 Flash resolved two opposite-effect groups and no all-enhancing groups. These counts partition the existing predictions; no additional model runs or changes to the reported overall accuracies are involved.

### A.5 Structural input processing

Inputs contained the mutation specification and a local structural neighbourhood extending 20 Å from the mutation site. Residue and atom identities were retained, while metadata, original identifiers and source annotations were withheld. Residue renumbering and record permutation reduced straightforward reconstruction of molecular identity without guaranteeing that identity was unrecoverable. Each complex was presented independently, without disclosing its mutation group or the effects measured for other partners. The same processed coordinate records were used with and without Python; Python and basic structural libraries could inspect distances, contacts and other structural relationships but did not supply dedicated energetic or mutation-effect predictions.

### A.6 Protein-identity recovery under input preprocessing

#### Task and inputs

This separate task asks the model to identify each protein chain’s family and specific protein name, not to predict mutation effects. For the processed-PDB condition, each attempt receives only /input/complex.pdb, /input/task.json and /input/instruction.txt. The task JSON contains only a task identifier and `task_type=protein_identity`; mutation fields, source names and answer labels are withheld. The loader does not further rewrite the supplied PDB bytes. It identifies protein chains using amino-acid C*α* records; non-protein components may provide context but are not identity targets. Reference labels, source-identity mappings and aliases remain on the host scoring side.

#### Model and analysis budget

The protocol specifies `deepseek-v4.1-flash` through an OpenAI-compatible interface, temperature 0.0, thinking enabled and `reasoning_effort=max`. Up to ten tool-assisted analysis rounds are allowed, followed by one tool-disabled JSON submission if no valid answer has been submitted. A valid earlier submission terminates the attempt. These are analysis rounds, not ten independent repetitions. The documented command uses the local Python runtime with eight workers and separate task messages and working directories.

#### Permitted analysis and scoring

The model may list and read files and run deterministic Python analyses of coordinates, contacts, composition, approximate secondary structure, disulfides and interfaces. Instructions prohibit internet or database access, literature or sequence/structure searches, other models and external prediction or simulation systems. Each chain must have one family hypothesis, one protein-name hypothesis, confidence and structural evidence; uncertain identities may be `null`. Host-side scoring accepts canonical labels or registered aliases after normalizing case, spaces and punctuation, without fuzzy substring matching. Family and protein-name accuracies are defined separately over chains with reference labels for each field. Null or wrong answers, incomplete chain coverage and invalid outputs are not counted as correct; unlabelled fields are excluded from the corresponding denominator. Repeated attempts are scored separately.

#### Reported aggregate results

The supplied summary gives 16/84 identity-recovery successes for processed, shuffled PDBs and 76/84 for the reference input (Table S2), a difference of 71.43 percentage points. These are identity-recovery counts, not mutation-effect group accuracies.

**Table S2:**
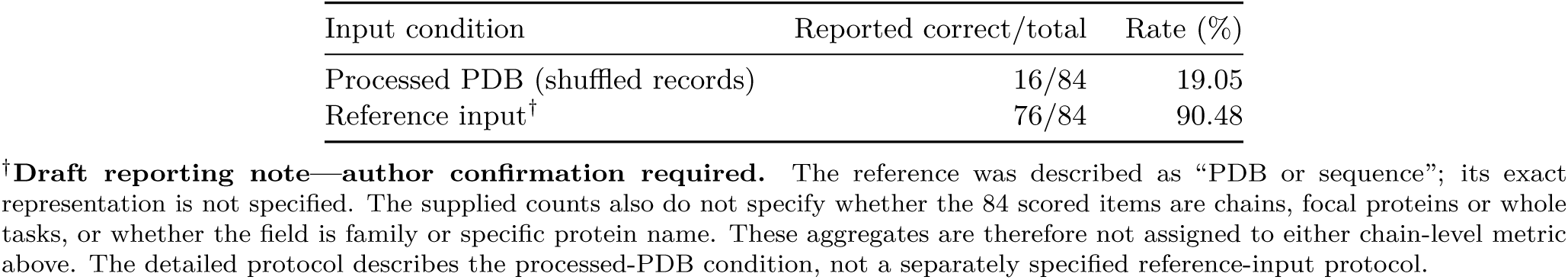
Reported aggregate identity-recovery results for DeepSeek Flash.

| Input condition | Reported correct/total | Rate (%) |
| --- | --- | --- |
| Processed PDB (shuffled records) | 16/84 | 19.05 |
| Reference input<sup>†</sup> | 76/84 | 90.48 |

<sup>†</sup>**Draft reporting note—author confirmation required.** The reference was described as “PDB or sequence”; its exact representation is not specified. The supplied counts also do not specify whether the 84 scored items are chains, focal proteins or whole tasks, or whether the field is family or specific protein name. These aggregates are therefore not assigned to either chain-level metric above. The detailed protocol describes the processed-PDB condition, not a separately specified reference-input protocol.

#### Interpretation

Subject to a common scoring definition across conditions, the contrast supports reduced identity recoverability after preprocessing, not elimination of memorization or proof of exclusively structural reasoning. It does not isolate the contributions of cropping, renumbering and record permutation. Family recognition and exact protein identification remain distinct.

## B Detailed case analyses

### B.1 LLMs capture partner-dependent effects from local structure

We first examined whether LLMs can recognize that the effect of a mutation depends on its local binding environment rather than simply on the intrinsic physicochemical similarity between the substituted amino acids. The ShPI-1 K13L case provides a direct test because the same P1 substitution produces opposite effects against two serine protease partners: K13L improves inhibition of bovine *α*-chymotrypsin by approximately 3.3-fold (PDB 3T62), but weakens inhibition of bovine cationic trypsin by approximately 118-fold (PDB 3M7Q) (Garćıa-Fernández et al., 2015).

GLM-5.3-Flash made correct predictions in both cases. For *α*-chymotrypsin, the model interpreted the wild-type (WT) lysine as a deeply buried charged group with limited electrostatic compensation (Fig. S1b). It reasoned that replacing lysine with leucine could both reduce the energetic cost associated with charge burial and maintain favorable packing within the specificity pocket. This interpretation is consistent with the experimental conclusion that increasing hydrophobicity at the P1 position is favorable in the relatively hydrophobic S1 pocket of *α*-chymotrypsin, as shown in Fig. S1b. The model also considered an alternative possibility—that the K*→*L substitution could eliminate favorable polar interactions—but ultimately assigned greater importance to the energetic gain from improved hydrophobic compatibility (Fig. S1e).

The same substitution was interpreted differently in the trypsin complex (Fig. S1c). In this case, GLM-5.3-Flash identified K13 as a deeply buried interfacial anchor stabilized by the acidic and polar environment of the S1 specificity pocket, including interactions with the Asp171/Ser172 region (Fig. S1d). The model therefore predicted that replacing lysine with leucine would disrupt this polar recognition network and that the shorter hydrophobic side chain would be unable to reproduce the interactions formed by the WT basic residue. This prediction is consistent with the experimentally observed substantial reduction in inhibitory activity.

We further constructed a decision tree based on nine reasoning trajectories generated by GLM-5.3, GLM-5.3-Flash, and DeepSeek-V4.1-Flash, as shown in Fig. S1e. The decision tree illustrates that the contrasting predictions made by the LLMs did not arise from an intrinsic rule that a K*→*L substitution is either favorable or unfavorable. Instead, the models evaluated the same chemical substitution within two distinct local interaction environments. These results demonstrate that, when the energetic consequences of a mutation are strongly determined by the surrounding binding-pocket environment, LLMs can capture structural determinants that depend on the specific binding partner.

A similar pattern was observed in the antibody case involving IgG1 Fc–Fc*γ*RIIa and IgG1 Fc–Fc*γ*RI (Ahmed et al., 2016), in which the same mutation increased affinity toward one receptor while decreasing affinity toward the other. The corresponding WT structures are shown in Fig. S2. Decision trees constructed from the reasoning trajectories further showed that the LLMs based their decisions on the distinct spatial configurations and geometric relationships present in the two receptor complexes (Fig. S2).

**Figure S1:**
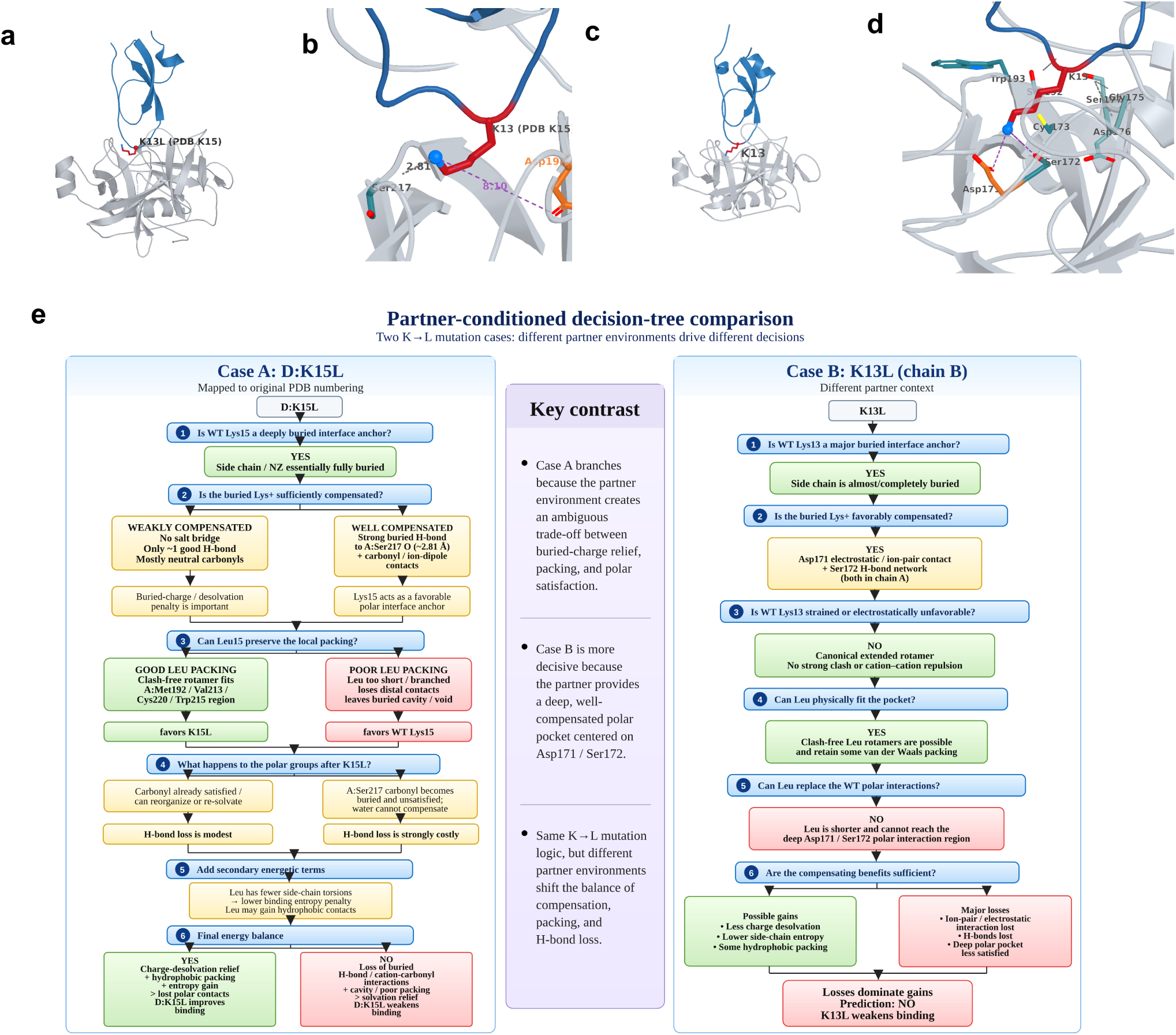
Structural analysis of ShPI-1 K13L in *α*-chymotrypsin and trypsin complexes. (a) Overall structure of the *α*-chymotrypsin–ShPI-1 complex; literature Lys13 corresponds to chain D Lys15 (PDB 3T62). (b) Lys15 is deeply buried (PDB 3T62). (c) Overall structure of the WT trypsin–ShPI-1 complex (PDB 3M7Q). (d) Lys13 forms a polar interaction network (PDB 3M7Q). (e) Decision tree summarized from the shared trajectories of three LLMs (GLM-5.3, GLM-5.3-Flash, and DeepSeek-V4.1-Flash).

**Figure S2:**
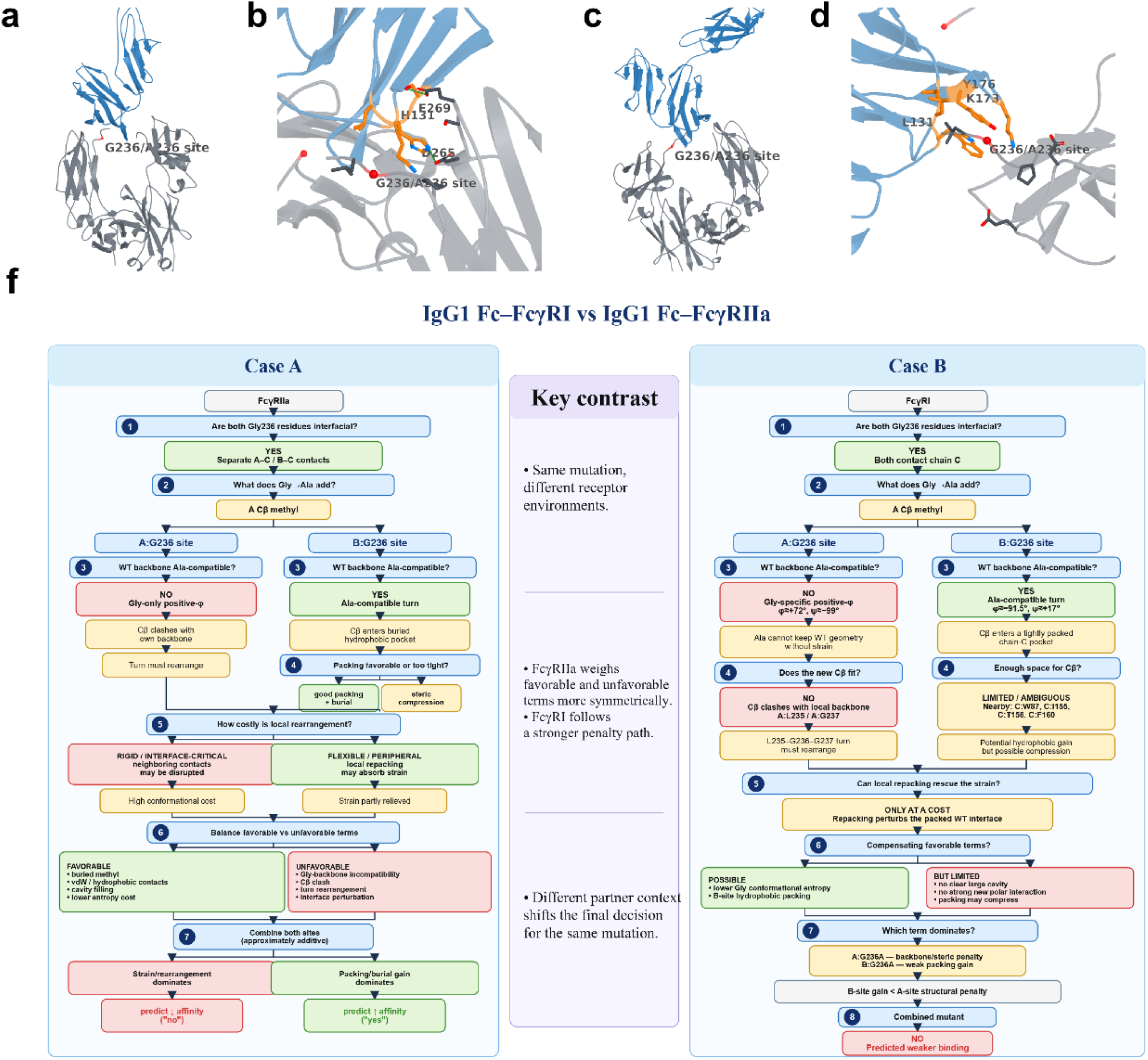
Structural context and representative LLM reasoning for the IgG1 Fc G236A mutation. (a) Overall structure of the IgG1 Fc–Fc*γ*RIIa complex (9MCX), highlighting the G236 mutation site. (b) Local structural environment surrounding the G236 site at the IgG1 Fc–Fc*γ*RIIa interface. (c) Overall structure of the IgG1 Fc–Fc*γ*RI complex (4ZNE), highlighting the G236 mutation site. (d) Local structural environment surrounding the G236 site at the IgG1 Fc–Fc*γ*RI interface. (f) Representative decision trees summarizing the LLM reasoning trajectories for the G236A mutation.

### B.2 Mutant-state hypotheses and energetic trade-offs

Next, we examined whether LLMs can infer plausible mutant-state interactions from wild-type (WT) structures and use these hypotheses to predict changes in binding effect. The BbKI F64R cases are particularly informative because experimentally determined mutant complex structures are available for direct comparison (Li et al., 2021). These include task 023db67447af92c088be, involving the WT BbKI–bovine trypsin complex (PDB 7JR1; mutant PDB 6DWH), and task 2530a32c128ad94f8dfc, involving the WT BbKI–KLK4 complex (PDB 7JQV; mutant PDB 7JOD).^1^

Taking the KLK4 complex as an example (Fig. S3a), GLM-5.3-Flash correctly predicted an increase in binding effect and proposed that Arg64 could compensate for the loss of Phe-mediated packing through interactions with the Asp189/Ser190 region and remodeling of the associated water network (Fig. S3b). Comparison with the mutant crystal structure partially supports this hypothesis. Arg64 indeed forms interactions with Ser190 and Asp189, indicating that the model identified the relevant region of interface remodeling. However, several specific contacts predicted by the model, including those involving Phe215 and Val227, were not observed in the experimental structure. In addition, the specific bidentate geometry proposed for the Arg64–Asp189 salt bridge is not established by the displayed hydrogen-bonding contacts alone (Fig. S3c). Thus, the model captured the overall trend of the interaction remodeling but did not accurately reconstruct the complete mutant interaction network. As illustrated by the decision tree, DeepSeek-V4.1-Flash proposed a similar hypothesis, suggesting that Arg64 could extend toward Asp189 and displace an ordered water molecule. Nevertheless, the model ultimately predicted decreased affinity because it assigned greater energetic weight to disruption of the tightly packed WT Phe environment.

**Figure S3:**
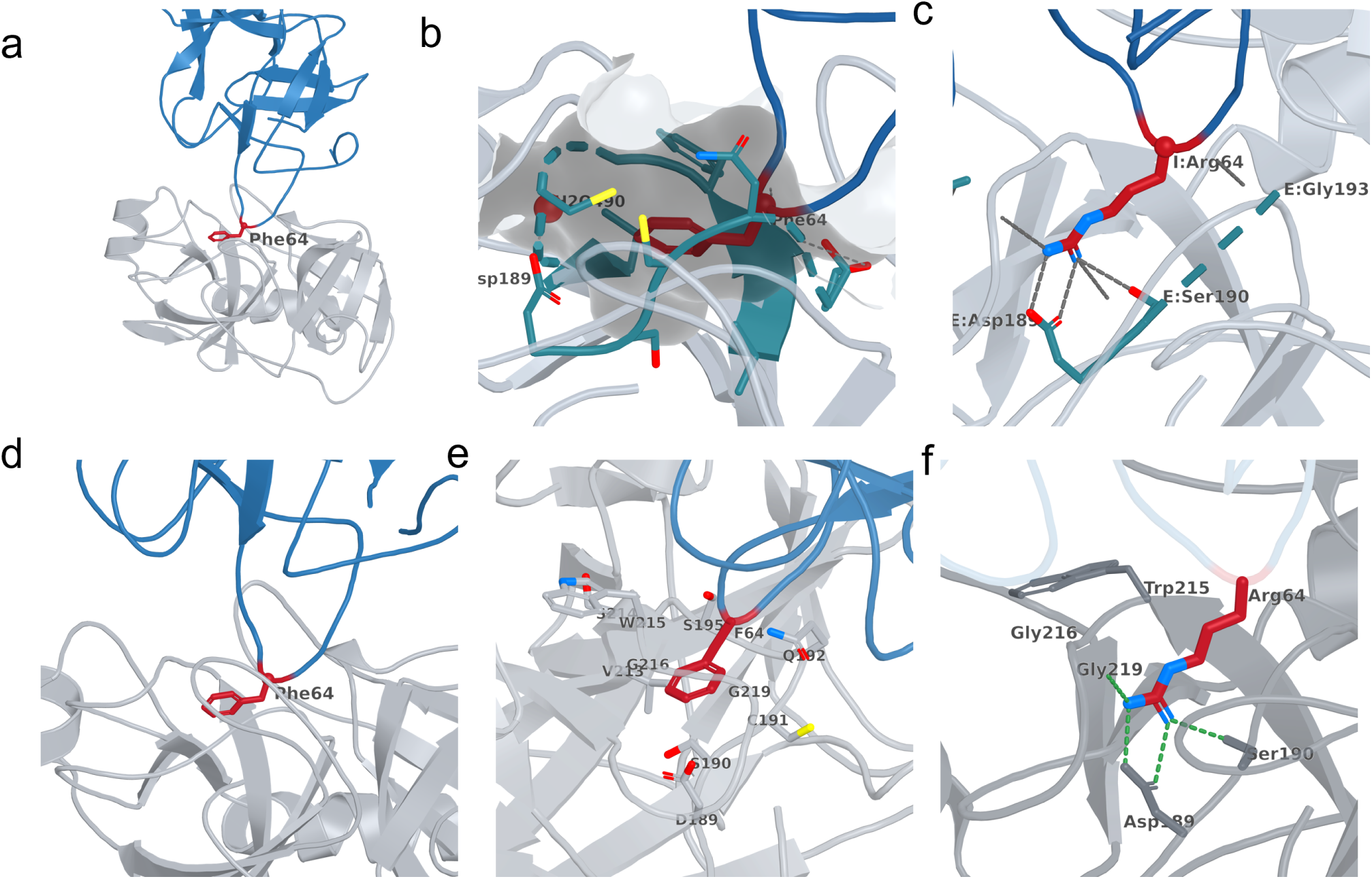
Structural analysis of BbKI F64R in KLK4 and trypsin complexes. (a) Overall structure of the 7JQV KLK4–BbKI complex, with Phe64 highlighted. (b) Local S1-pocket environment around Phe64; Asp189 lies at the pocket bottom, with HOH490 nearby. (c) Crystallographically observed interaction pattern of Arg64 in the 7JOD mutant complex. (d) Overall structure of the BbKI–trypsin complex, with residue 64 highlighted at the interface. (e) Local environment of WT Phe64 within the tightly packed trypsin interface pocket. (f) In the mutant complex, Arg64 forms hydrogen-bonding interactions with Asp189, Ser190, and Gly219.

A similar uncertainty was observed in the second F64R complex (WT: 7JR1; mutant: 6DWH; Fig. S3d). Both the “yes” and “no” reasoning trajectories identified the key role of Asp189 and the possibility of newly formed interactions. Their main disagreement concerned energetic weighting: the “yes” trajectories considered the enhanced polar interactions mediated by Arg64 sufficient to favor binding, whereas the “no” trajectories judged these gains insufficient to compensate for the loss of hydrophobic packing contributed by F64 (Fig. S3e). We further examined the mutant-state interactions predicted by the LLMs. As shown in the figure, the models hypothesized that Arg64 could form hydrogen bonds with Ser190, Gly219, and Gly216. The crystal structure confirmed hydrogen-bonding interactions with Gly219 and Ser190, but not with Gly216. Arg64 also formed a hydrogen-bonding interaction with Asp189; this description alone does not exclude a salt bridge (Fig. S3f). In addition, all models made unfavorable predictions for the corresponding mutation at the same site in the BbKI F64R–*α*-chymotrypsin complex (PDB 7JRX), further illustrating the ability of LLMs to discriminate mutation effects according to the local structural environment (Fig. S4).

These results further indicate that, when LLMs make judgments based on rational physicochemical knowledge, their uncertainty mainly arises from two stages: first, extrapolating unobserved mutant conformations from the observed WT structure; and second, translating local changes in interactions into changes in the overall binding free energy. Reasoning trajectories that produced favorable predictions placed greater emphasis on the possibility that Arg64 could form new interactions with the buried acidic site, whereas trajectories that produced unfavorable predictions placed greater emphasis on the loss of hydrophobic and aromatic packing present in the WT complex. These findings indicate that LLMs can generate chemically plausible hypotheses about mutant-state interactions from WT coordinates, but remain inconsistent in determining whether newly formed interactions can compensate for the loss of pre-existing interactions. Their principal limitation therefore lies not only in inferring the local structure after mutation, but also in appropriately weighting these competing energetic contributions.

### B.3 WT-state evidence limits inference of indirect interface remodeling

Next, we used the EVA-A–CCL7 cases (Devkota et al., 2023) to illustrate another limitation of LLM reasoning: models can generate mechanistic hypotheses that are consistent with experimentally observed mutation effects, yet fail to assign these hypotheses sufficient weight when making the final prediction. We focused on two affinity-enhancing mutations, Y44A (WT: 7SCU; mutant: 8FK6; Fig. S5a) and L39P (WT: 7SCU; mutant: 8FK8; Fig. S5d), both of which have experimentally determined mutant complex structures and are located at the interface. In both cases, although the models accurately described key features of the wild-type (WT) binding pocket, they predicted that the mutations would not improve affinity, in disagreement with the experimental outcomes.

Further analysis of the reasoning trajectories showed that, for the EVA-A Y44A–CCL7 mutation (task ID: task 85c20dee617ba09a64f1), the models identified Tyr44 as being located within the pocket surrounding CCL7 Tyr13 (Fig. S5b). Its aromatic side chain is largely buried within EVA-A and participates in local packing, which ultimately led the models to infer that the Y44A mutation would remove favorable interactions and weaken the stability of the binding interface. However, the mutant structure 8FK6 and the published study show that the affinity enhancement does not arise from new direct interactions contributed by Ala44 (Fig. S5c). Instead, removal of the bulky Tyr side chain changes the conformational requirements of the pocket and reduces the structural rearrangement required to accommodate CCL7 Tyr13. Importantly, this possibility was not entirely absent from the model reasoning. The reasoning trajectories explicitly considered a favorable counterfactual scenario in which removal of Tyr44 could open the pocket and allow the partner Tyr residue to adopt a lower-strain and more tightly packed conformation. Nevertheless, this hypothesis was ultimately rejected by LLMs because the observed WT complex already exhibited a tightly packed interaction network and showed no obvious steric clashes.

The EVA-A L39P–CCL7 case (task ID: task 01d69c7d396248df43b2) provides an even more representative example. Based on the WT complex, the models identified Leu39 as contributing to local hydrophobic packing (Fig. S5e). DeepSeek-V4.1-Flash and GLM-5.3-Flash therefore expected that replacing Leu39 with proline would disrupt favorable packing interactions, eliminate a backbone hydrogen-bond donor, and would not improve affinity. However, both the published study and the mutant structure 8FK8 show that this rearrangement is critical for the affinity enhancement. Pro39 reshapes the local pocket, allowing CCL7 Tyr13 to insert more deeply into the CC+1 binding site and form more favorable aromatic interactions with EVA-A Tyr44 and His28 (Fig. S5f). Similar to the Y44A case, the affinity enhancement does not arise from a new favorable interaction contributed directly by the mutated side chain, but instead emerges indirectly through interface remodeling.

The model trajectories again show that this mechanism was, to some extent, anticipated. DeepSeek-V4.1-Flash explicitly considered that removal of the space occupied by the Leu side chain could “allow the Tyr to relax into the void, possibly making better contacts,” whereas GLM-5.3-Flash considered whether a “Pro-induced kink could reposition” the interface to improve contacts. Although these hypotheses are not identical to the experimentally observed mechanism, they capture its central feature: mutation-induced remodeling allows the ligand to adopt a more favorable conformation. However, both models assigned low confidence to these hypotheses because the structural relaxation they anticipated extended beyond the geometry observed in the WT structure. In contrast, the models placed greater emphasis on directly observable changes, namely the loss of tight packing and hydrogen-bonding interactions present in the WT structure.

**Figure S4:**
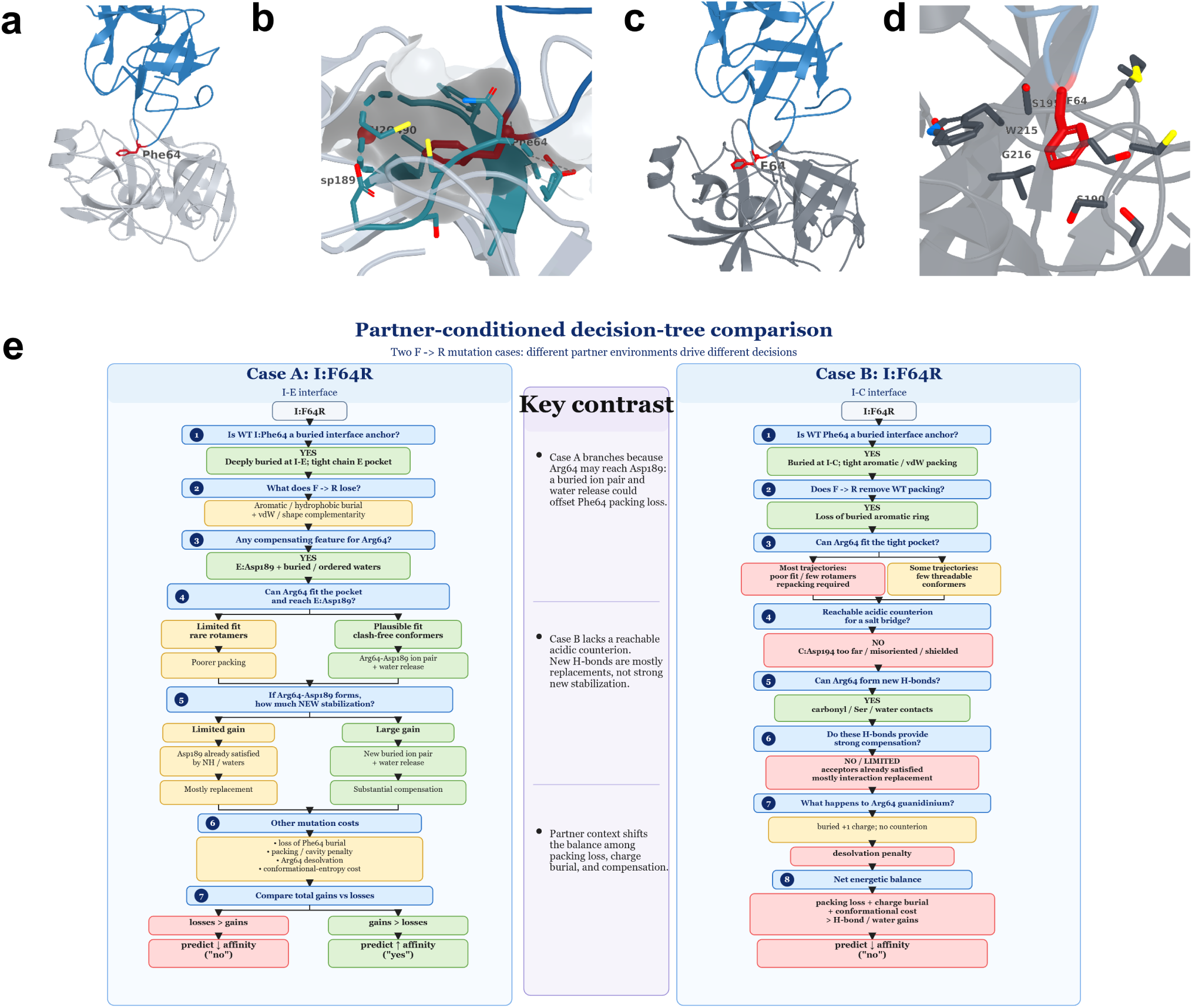
Structural analysis and LLM reasoning for the BbKI F64R mutation across different protease partners. (a) Overall structure of the BbKI–KLK4 complex (PDB 7JQV). (b) Local structural environment surrounding BbKI Phe64 in the KLK4 complex. (c) Overall structure of the BbKI–*α*-chymotrypsin complex (PDB 7JRX). (d) Local structural environment surrounding BbKI Phe64 in the *α*-chymotrypsin complex. (e) Representative decision trees summarizing the LLM reasoning trajectories for the F64R mutation in the two partner contexts, with the BbKI–KLK4 complex shown on the left and the BbKI–*α*-chymotrypsin complex shown on the right.

**Figure S5:**
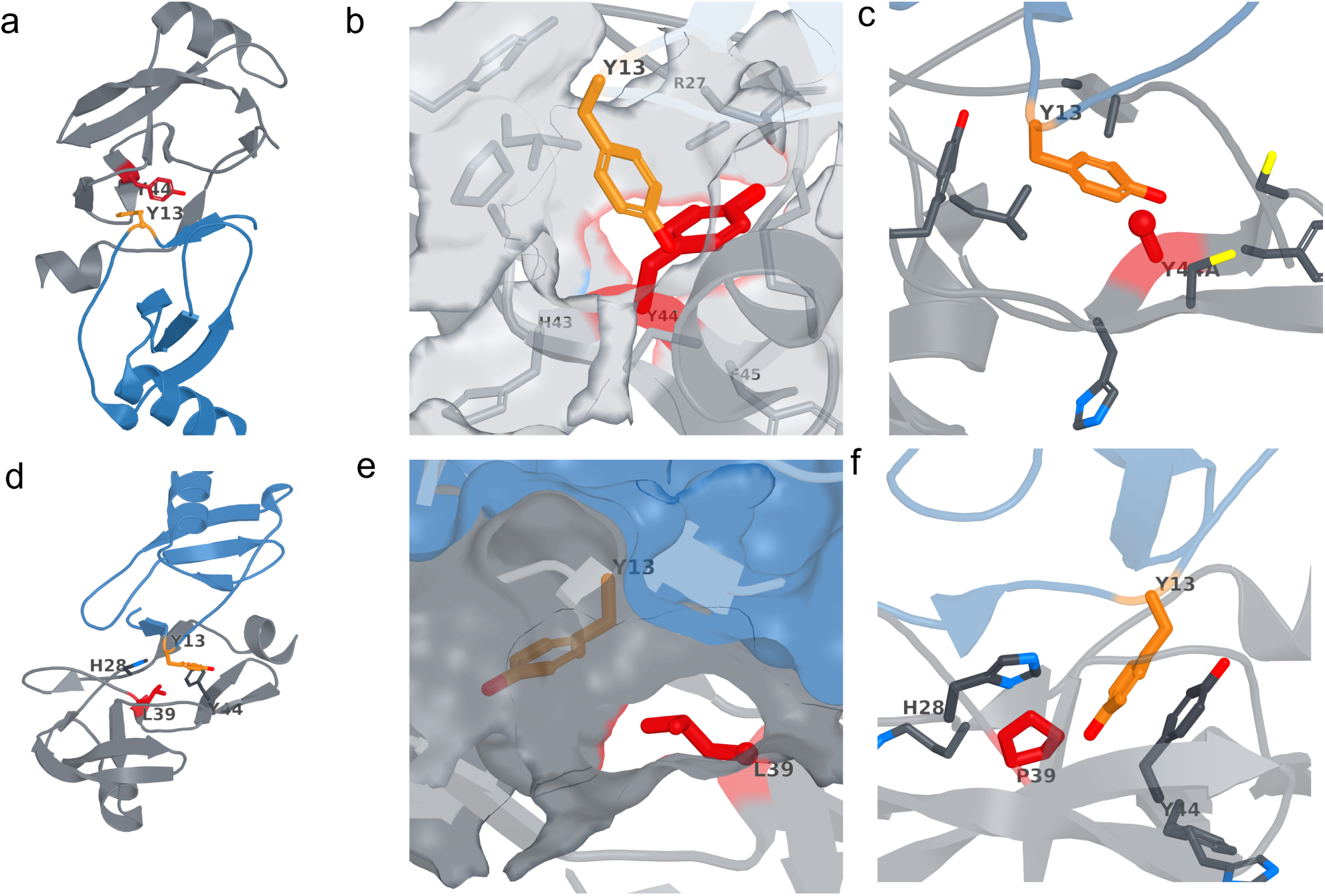
Structural analysis of EVA-A Y44A and L39P in the EVA-A–CCL7 complex. (a) Overall WT EVA-A–CCL7 complex (PDB 7SCU), with EVA-A Tyr44 and CCL7 Tyr13 highlighted. (b) Local structural environment of EVA-A Tyr44 in the WT complex. (c) Y44A mutant complex (PDB 8FK6), showing the remodeled interface after removal of the Tyr44 side chain. (d) Overall WT EVA-A–CCL7 complex used for analysis of L39P. (e) Local WT interface around EVA-A Leu39 and CCL7 Tyr13. (f) L39P mutant complex (PDB 8FK8), showing the altered interface configuration induced by Pro39.

### B.4 Correct prediction with incorrect reasoning

Finally, we used the BLIP E73W case (Rivera et al., 2025) to illustrate a distinct failure mode in which LLMs predict the correct direction of the mutation effect but attribute it to an incorrect or incomplete structural mechanism. Although GLM-5.3 Flash correctly predicted the affinity enhancement and identified several valid wild-type (WT) structural features (Fig. S6a and Fig. S6b), the proposed mechanistic explanation was not consistent with the mechanism described in the literature. The model attributed the affinity improvement mainly to the removal of the unfavorable energetic penalty associated with the buried Glu residue, water displacement, and aromatic repacking. However, the mutant structure revealed a distinct interaction mode involving packing between Trp73 and Tyr105 and stabilization of an alternative partner conformation (Fig. S6c and Fig. S6d). This case shares a similar failure mode with the EVA-A cases (Y44A and L39P). When LLMs lack relevant prior knowledge or the ability to reason beyond static geometric information, they face challenges in more complex mutation-effect interpretation problems.

**Figure S6:**
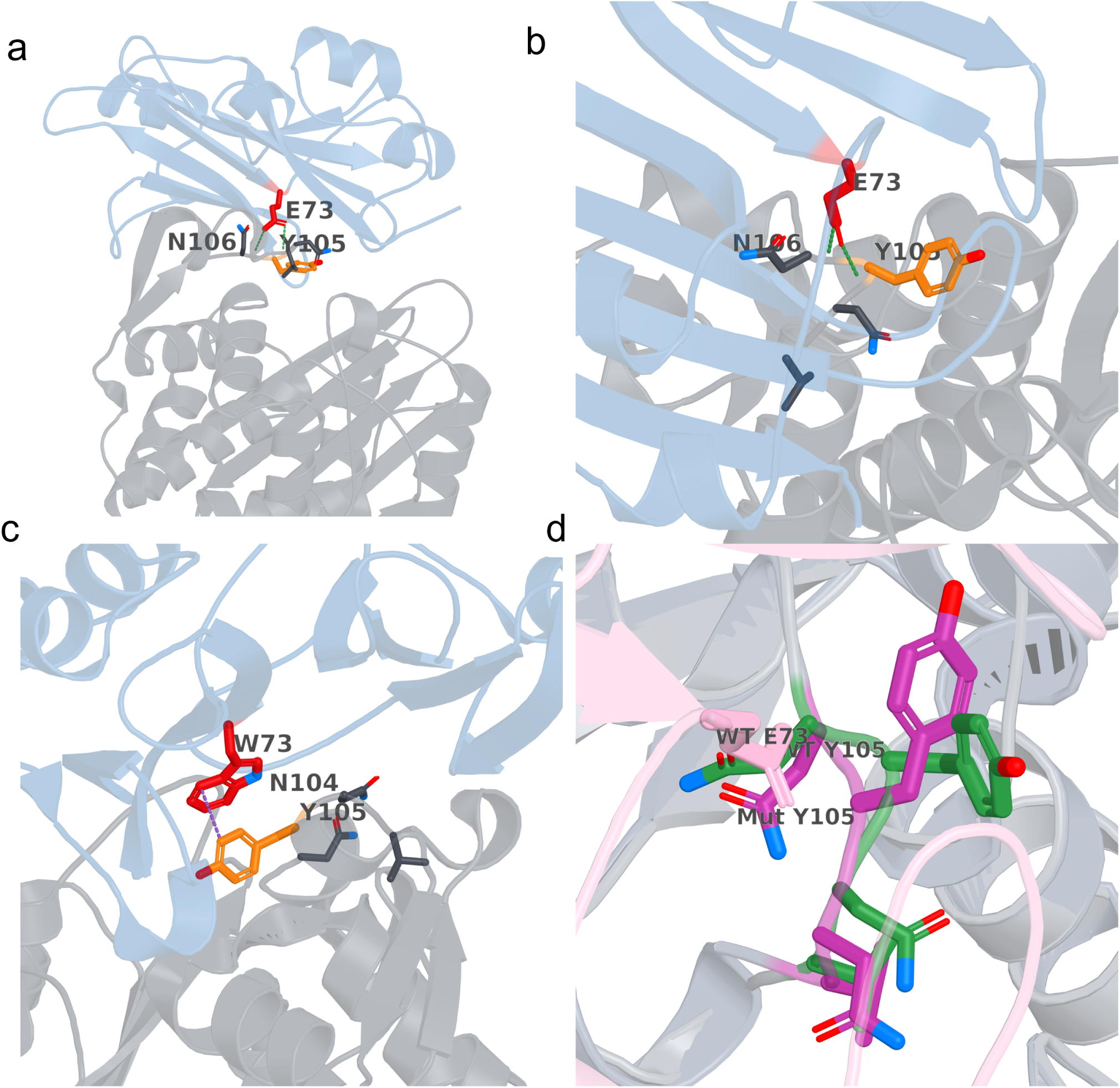
Structural analysis of BLIP E73W in the BLIP–CTX-M-15 complex. (a) WT BLIP–CTX-M-15 complex (PDB 7S5S). (b) Local WT interface around BLIP Glu73 and the CTX-M-15 103–106 loop. (c) E73W mutant complex (PDB 9Q0B), showing the remodeled local interface. (d) Structural overlay of WT and E73W complexes, highlighting rearrangement of the 103–106 loop and Tyr105 around BLIP residue 73.

## C Protein-system distribution of successful judgments

Figure S7 gives the system-level counts underlying Section 3.4. All LLM rows are Flash models. The Python-assisted DeepSeek, GLM and Qwen3.8 rows cover four, five and four systems, respectively, and collectively seven of the 17 Dataset 1 systems. Counts are reproduced from the supplied system summary; they do not identify exact individual-group overlap or add independent model runs.

**Figure S7:**
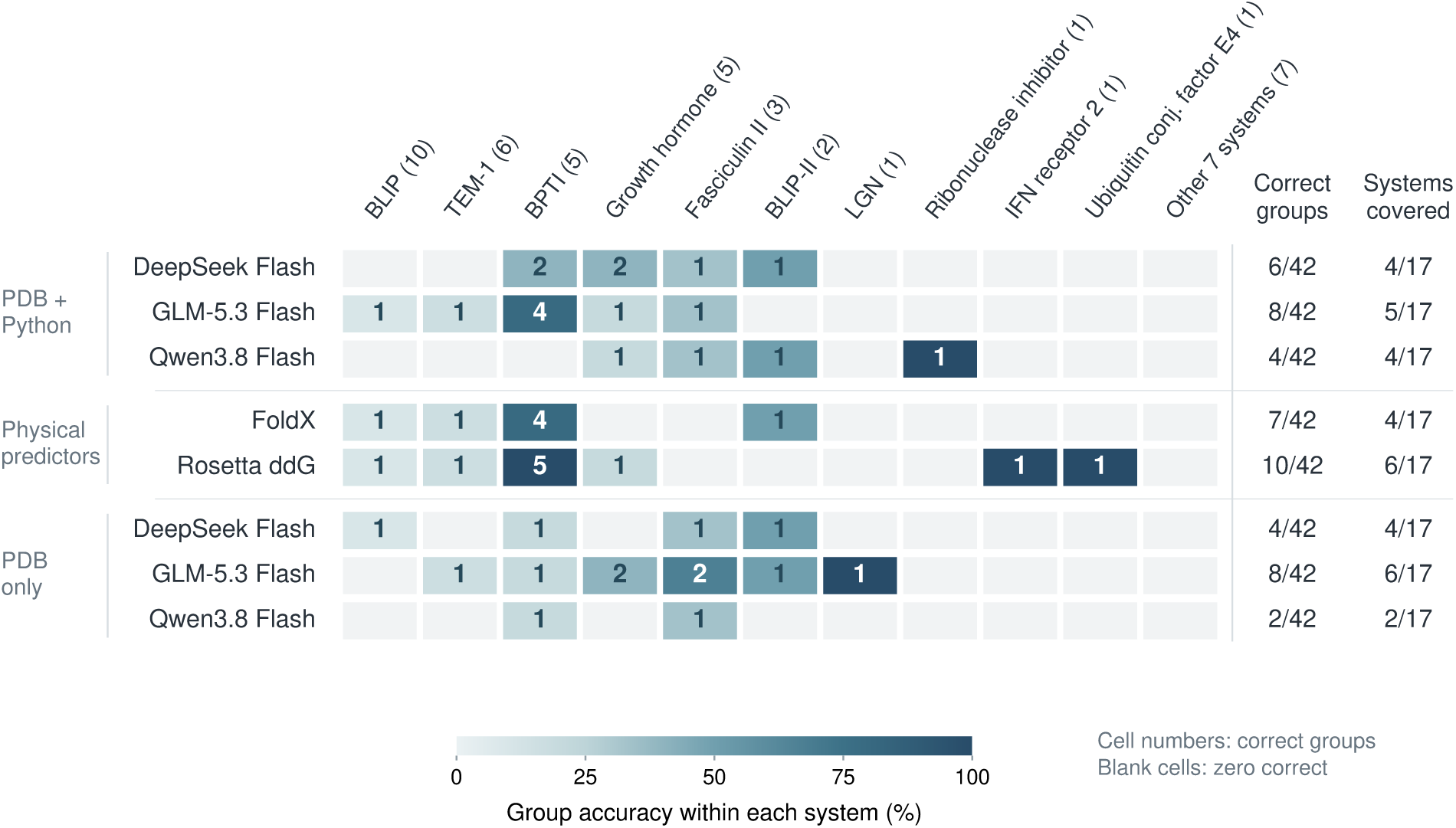
Successful judgments across protein systems. Cells show correct-group counts; shading denotes within-system group accuracy on Dataset 1. Blanks indicate zero. Column-label parentheses give group counts; the seven pooled systems have no successes. Right columns report correct groups and systems covered. All LLM rows are Flash models. System-level counts do not establish individual-group overlap.

## D Structural-claim extraction and audit procedure

This appendix details the structural-claim audit in Section 4.3. Mol5 refers here to the Mol* 5.11.0 interaction analysis.

### D.1 Claim extraction and audit scope

We scanned the structural-evidence text in each prediction.json file and split independently verifiable relationships within a sentence into atom- or residue-level claims. For each claim, we recorded the mutated residue, interaction type, whether the relationship was asserted or negated, interaction target, chain, residue number, reported distance, inter-chain or intra-chain scope, and backbone or side-chain scope. A single evidence entry could yield multiple claims; evidence-entry counts and claim counts therefore need not be equal.

The recognized interaction types were hydrogen bonds, weak hydrogen bonds, ionic interactions or salt bridges, water bridges, hydrophobic contacts, *π*–*π* stacking, cation–*π* interactions, halogen bonds and metal coordination. Solvent-accessible surface area (SASA), cavity volume, burial, B factors, flexibility and hypothesized post-mutation rearrangements were not directly mapped to Mol5 interaction claims and were outside the audit scope.

### D.2 Structure analysis and claim matching

Input PDB structures were analyzed with Mol* 5.11.0, with all interaction providers enabled and the default Mol5 distance and geometry criteria retained. Claims were matched to Mol5 records by task, PDB, mutation site, interaction type, target chain and residue. Where the model reported a distance, a numerical tolerance of 0.15 Å was applied only when the distance definitions were consistent.

### D.3 Manual review

All 12 records automatically flagged as conflicts were subsequently inspected manually. Particular attention was given to records involving negation, multiple residues, multiple distances, water bridges, salt-bridge distances, and ambiguity about backbone versus side-chain scope.

## Footnotes

1 For these BbKI tasks, WT and mutant designate the supplied Phe64 starting state and the Arg64 comparison state, respectively. The Phe64 structures are R64F variants of natural BbKI; the prediction task is the reverse F64R substitution.

